# Oxytocin and vasopressin 1a receptor alterations in the superior temporal sulcus and hypothalamus in schizophrenia

**DOI:** 10.1101/2024.08.23.609421

**Authors:** Ariel W. Snowden, Sarah E. Schwartz, Aaron L. Smith, Mark M. Goodman, Sara M. Freeman

## Abstract

Schizophrenia is a chronic, severe psychiatric condition characterized in part by social impairments. As social cognitive functioning is a major predictor of successful life outcomes in schizophrenia, there is a critical need to determine the neurobiological basis of social impairments in schizophrenia. The current study used receptor autoradiography to examine vasopressin 1a (AVPR1a) and oxytocin receptor (OXTR) densities in the postmortem brain tissue of individuals who had schizophrenia (N=23) relative to unaffected matched controls (N=18). We analyzed the superior temporal sulcus, a brain region that is highly implicated in social perception and shows aberrant functioning in schizophrenia. GreaterAVPR1a binding densities were observed relative to OXTR. Increases in AVPR1a densities with advanced age in females who had schizophrenia were also observed. This finding in the superior temporal sulcus may explain a shift in positive symptom severity(paranoia/suspiciousness) that has been observed with advanced age in women with schizophrenia. In addition, increased OXTR and a trend toward increased AVPR1a densities were observed in the hypothalamus in schizophrenia. The hypothalamus synthesizes oxytocin and vasopressin in the brain and is the initiator of the hypothalamic-pituitary-adrenal axis, which facilitates the fight or flight stress response. These receptor systems may be upregulated to compensate for possibly low exogenous oxytocin and vasopressin release from the hypothalamus in schizophrenia. Our findings show age- and sex-dependent effects on nonapeptide receptor binding that shed light onto the neurobiology of the social brain in the progression of schizophrenia.

## 1. Introduction

Schizophrenia (SZ) is a severe, chronic disorder affecting about 2 million individuals in the United States alone^1^. SZ is characterized by hallucinations, delusions, cognitive deficits, and impairments in social cognition. Although antipsychotic drugs are typically effective in treating hallucinations and delusions in SZ, the social cognitive symptoms (e.g., social withdrawal, impaired emotion perception) often persist or are exacerbated by these treatments^2^. As social cognitive functioning predicts successful life outcomes in SZ^2^, there is a critical need to determine the biological basis of social cognitive impairments in SZ.

Due to the ability of the neurohormones oxytocin (OXT) and vasopressin (AVP) to facilitate many social behaviors in animals^3^ and humans^4^, variation in central OXT and AVP may underlie social cognitive impairments in SZ. In neurotypical humans, intranasal OXT (IN-OXT) affects several domains of social cognition, including eye gaze toward social stimuli, emotion recognition, and trust^4^. AVP is structurally similar to OXT, differing at just two positions in the nine-amino acid chain, and is implicated in territoriality, aggression, and pair bonding^5^. Intranasal AVP (IN-AVP) administered to nonclinical human participants affects social cooperation, social attention, and face processing^6^.

Many studies have examined potential variation in AVP and OXT in SZ using plasma hormone concentrations. Elevated plasma OXT levels, reduced plasma OXT, and no differences in plasma OXT^7^have been reported in SZ relative to unaffected controls. Although baseline OXT levels between SZ and controls are inconsistent, reduced OXT levels are often associated with more severe negative symptoms (e.g., social withdrawal^7^). Peripheral AVP studies have also yielded mixed results, observing higher AVP levels^8^, reduced AVP^9^ and no differences in plasma AVP^10^ between SZ and controls. Interestingly, elevated AVP plasma concentrations in SZ relative to controls are observed during periods of high psychological stress^8^. In particular, increased plasma AVP levels in women with first-episode psychosis are linked to worse positive symptoms and higher suspiciousness^8^. These studies suggest that variations in endogenous OXT and AVP levels in SZ contribute to social cognitive symptoms. However, there are several methodological issues in measuring plasma OXT and AVP: (1) Peptide assays may bind to nonspecific molecules with a higher affinity than the target hormone, resulting in inflated concentrations^11^; (2) Baseline plasma concentrations are highly variable, especially when measuring extracted OXT plasma levels^11^; (3) Plasma levels are often not correlated with cerebrospinal fluid levels, indicating that plasma measures cannot reliably elucidate central OXT and AVP levels^12^.

Due to the aforementioned inconsistencies in findings from plasma studies, there is a critical need to examine the OXT and AVP systems directly in the human brain. Until recently, isolating OXTR binding from AVPR1a binding in postmortem primate and human brain tissue was challenging due to the structural similarity of these receptors^13^. Both OXT and AVP, along with their radioligands, have high affinities for both OXTR and AVPR1a, complicating the accurate identification of OXTR location^13^. Consequently, early reports of OXTR distribution in the human brain have been considered inconclusive because of the confounding AVPR1a signal^14^. These same challenges are observed in the only known study examining OXTR in postmortem brains of individuals who had SZ. Uhrig and colleagues^15^ report decreased cerebellar vermis OXTR and decreased *OXTR* mRNA in the temporal cortex in SZ. The authors conclude that there is downregulated OXTR within the social cognitive network in SZ. However, due to the mixed affinity of the OXTR radioligand for both the OXTR and AVPR1a, their findings are likely confounded by overlapping AVPR1 binding and thus, are inconclusive.

To address these limitations, we used a previously validated receptor autoradiography technique^13^ to isolate OXTR and AVPR1a in human brain tissue from donors with or without SZ. To this end, the current study characterized OXTR and AVPR1a binding densities in regions affected during social perception in SZ (superior temporal sulcus; STS) and involved in OXT and AVP synthesis (hypothalamus).

The STS is a cortical region of the temporal lobe that is highly implicated in social perception, including face evaluation, speech perception, theory of mind (interpreting others’ mental states), and audiovisual integration^16^. In SZ, the STS is often hyperactive during emotionally neutral face perception, which is associated with a risk allele (rs1334706) most strongly linked to a SZ diagnosis^17^. STS hypoactivation during social processing, such as theory of mind tasks, is also reported in SZ^18^. Notably, IN-OXT appears to increase STS activation, which is associated with improved accuracy during theory of mind tasks^18^. Furthermore, increased resting state amygdala-STS functional connectivity is observed with IN-OXT administration in SZ^19^.These findings suggest that IN-OXT recruits brain regions that are typically disengaged during social perception in SZ. However, the mechanisms by which IN-OXT exerts its effects in the STS are unknown. We hypothesized that STS OXTR densities would be lower in SZ compared to controls, potentially contributing to social cognitive impairments and the inability to recruit the STS during complex social tasks in SZ. Additionally, we expected higher STS AVPR1a binding densities in SZ, potentially related to hypervigilance and heightened threat perception during neutral face processing in SZ.

The hypothalamus regulates the sleep-wake cycle, the stress response, and synthesis and secretion of hormones^20^. Enlarged hypothalamic volumes are observed in individuals with SZ and in their first-degree relatives, compared to unaffected controls^21^. Further, individuals with SZ show decreased AVPR1a mRNA expression within the paraventricular nucleus^22^, a subregion of the hypothalamus that synthesizes OXT and AVP^4^. In animals, the paraventricular nucleus of the hypothalamus is involved in maternal aggression and avoidance behaviors in response to stressful events^3^. We hypothesized that there would be increased hypothalamic OXTR in SZ compared to controls. Individuals with SZ typically exhibit elevated baseline cortisol levels, suggesting overactivity of the hypothalamic-pituitary-adrenal axis^23^. OXT has been shown to reduce stress response by regulating this axis in rodents and primates^24^. Therefore, reduced OXT levels in the hypothalamus could lead to both increased cortisol levels and upregulated OXTR. Our findings shed light on both OXT and AVP functioning within the hypothalamus and STS in SZ.

## 2. Methods

### 2.1 Specimens

Unfixed, frozen blocks of de-identified, postmortem human brain tissue were obtained from University of Miami Brain Endowment Bank, University of Maryland Brain and Tissue Bank, Harvard Brain Tissue Resource Center, The Human Brain and Spinal Fluid Resource Center and Mt. Sinai Brain Bank, which are Brain and Tissue Repositories of the NIH NeuroBioBank. Information about postmortem tissue processing is detailed on the NeuroBioBank website (https://neurobiobank.nih.gov/). An *a priori* power analysis was conducted in G Power, indicating that a total sample size of 42 (21 per group, with 3 repeated-measures per subject) is sufficient to obtain a statistical power of 95% for a mixed design analysis of variance (MIX-ANOVA) with a within-between interaction. Although a total of 42 specimens were initially requested from each brain region from the NIH Neurbiobank, the total number of specimens per region varied based on the actual specimens available in the NeuroBioBank. STS specimens received were from a total of 41 human donors: 23 SZ (12 female, 11 male) and 18 unaffected controls (8 female, 10 male). The mean age of donors did not significantly differ across groups (64.3±9.6 in the SZ group and 68.0±10.9 in the unaffected control group) for the STS. Hypothalamus specimens received were from a total of 28 human donors: 14 SZ (6 female, 8 male) and 14 unaffected controls (6 female, 8 male). The mean age of donors did not significantly differ across groups (66.21± 11.70 in the SZ group and 67.21 ± 10.21 in the unaffected control group) for the hypothalamus. Inclusion criteria for SZ specimens was a diagnosis of SZ (unspecified, undifferentiated, residual, or paranoid types). Inclusion criteria for control tissues included cases matched as best as possible on age, race, and sex to the SZ cases. Demographic information and additional donor information is provided in Supplementary Table 1. Exclusion criteria for SZ donor samples were any comorbidities for other clinical brain diagnoses (e.g., epilepsy, Parkinson’s disease, autism spectrum disorder, a diagnosis of catatonic SZ, comorbidities for other psychiatric diagnoses (e.g., bipolar disorder), intellectual disabilities, and any cause of death other than natural (e.g., suicide). Exclusion criteria for matched unaffected control samples were any history of psychiatric diagnoses, intellectual disabilities, and any cause of death other than natural.

### 2.2 Tissue preparation and anatomical details

All specimens were shipped on dry ice and immediately stored at −80°C from receipt until sectioning. Brain blocks were sectioned coronally at 20 µm on a cryostat at −15-20°C and thaw-mounted to Superfrost-Plus slides. Slides were then stored at −80°C in a sealed slide box with a desiccant packet until use in assays. The specimens showed slight individual anatomical variations within Brodman’s area 22. We used the Allen Human Brain Atlas (https://atlas.brain-map.org/) to identify the STS and the anatomical boundaries of the gyri encompassing the sulci: superior temporal gyrus and the middle temporal gyrus (inferior to the STS); see Figure 1. Due to anatomical variation in the location of the hypothalamic blocks and the fact that each block was removed from gross neuroanatomical landmarks, we were unable to confidently determine the identities of any subnuclei (e.g., paraventricular and supraoptic nuclei) where binding was measured.

**Figure 1.**
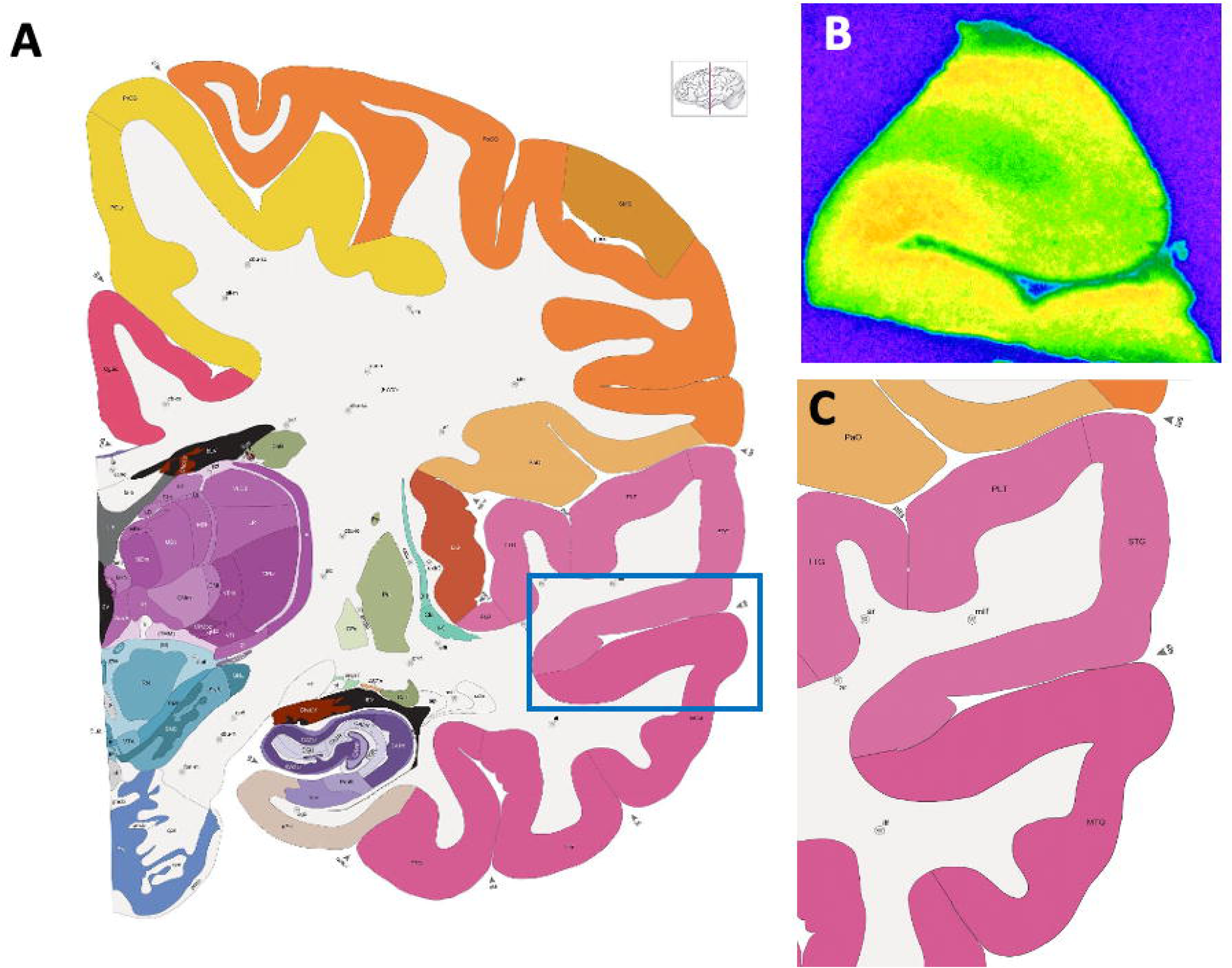
Allen Human Brain Reference Atlas Showing the Anatomy of the Superior Temporal Gyrus. A) Whole hemisphere coronal section showing the area quantified (within the blue box); B) Pseudocolored autoradiogram displaying AVPR1a receptor binding in the superior temporal gyrus extending to the middle temporal gyrus; C) A closer examination of the region encompassing the superior temporal sulcus (STS), which includes the superior temporal gyrus (STG) and the middle temporal gyrus (MTG). Atlas images (A, C) are available under the Creative Commons License Attribution-NonCommercial-NoDerivs License and were originally published in^50^.

### 2.3 Competitive binding receptor autoradiography

Slides were processed for OXTR and AVPR1a competitive binding receptor autoradiography according to detailed methods previously published. Briefly, sections were thawed in a sealed slide box containing a desiccant packet for 1 hour at room temperature. After a light fixation in 0.1% paraformaldehyde (pH 7.4) and washes in Tris Buffer, adjacent series of sections were incubated with either 90 pM of the OXTR radioligand, ^125^I-ornithine vasotocin analog (^125^I-OVTA) or 30 pM of the AVPR1a radioligand, ^125^I-linearized vasopressin antagonist (^125^I-LVA). These concentrations were determined to be at the radioligand’s binding affinity for the appropriate human receptor^13^. Adjacent sections for each of the two radioligands were incubated in the following three conditions: 1) radioligand alone, 2) radioligand plus 1 nM of the selective AVPR1a antagonist SR49059, or 3) radioligand plus 100 nM of the novel OXTR antagonist, ALS-II-69. SR49059 is available from Tocris (Minneapolis, MN) and has been shown *in vitro* to selectively occupy human AVPR1a over human OXTR in the 1-10 nM range^13^. ALS-II-69 is a human-selective OXTR antagonist that is highly selective for the human OXTR at concentrations up to 1 µM^13^. After radioligand incubation (with or without selective competitors), the slides were washed with Tris buffer, dipped in deionized water, and allowed to air dry. Finally, slides were exposed to radiographic film: films from the OXTR assay were exposed to Carestream Biomax XAR film (Kodak, Rochester, NY) for 5 days. Slides from the AVPR1a assay were exposed to Ultracruz Blue Autoradiography films (Santa Cruz Biotechnology, Inc., Dallas, TX) for 8 days. This change in film type between the OXTR and AVPR1a assays was due to the discontinuation of the XAR films before the AVPR1a assay was completed. All films were exposed to a set of ten ^125^I standard microscales (American Radiolabeled Chemicals, St. Louis, MO) then developed and analyzed.

### 2.4 Quantification of film autoradiograms

Optical binding density (OBD) was quantified from the film autoradiograms using the MCID Digital Densitometry Core System (Interfocus Imaging, Cambridge, UK). Images of the film were loaded into the software using a light box and top-mounted camera connected to a computer. A flat-field correction for luminosity levels was equally applied to all images. OBD values from the set of autoradiography standards were then loaded into the software and used to generate a standard curve, from which OBD values were extrapolated. Experimenters were blind to the diagnostic condition during quantification of the films.

#### 2.4.1 Hypothalamus

For each specimen, three OBD values were measured according to the boundaries of the binding density. An average background OBD value was subtracted from this average hypothalamus measurement in order to account for any individual differences in nonspecific binding and to yield normalized OBDs across specimens. Determination of the precise subnuclei of the hypothalamus was not possible given the anatomical variation from specimen to specimen and the lack of other neuroanatomical landmarks.

#### 2.4.2 Superior Temporal Sulcus

We utilized the competitive binding approach to account for any individual variation in nonspecific binding in the following way. For the OXTR analysis, there are three conditions: A) OXTR radioligand alone (resulting signal includes both OXTR and AVPR1a binding); B) OXTR radioligand + OXTR competitor (should reveal any AVPR1a binding); C) OXTR radioligand + AVPR1a competitor (should reveal the OXTR signal). For the AVPR1a analysis, there are three conditions: D) AVPR1a radioligand alone (resulting signal included both AVPR1a and OXTR binding); E) AVPR1a radioligand + AVPR1a competitor (should reveal any OXTR binding), and F) AVPR1a radioligand + OXTR competitor (should reveal only AVPR1a signal). Due to a lack of consistent background area in the blocks of cortical tissue for the STS, we used an alternative condition subtraction approach to account for any individual variation in nonspecific binding. For the OXTR radioligand, we subtracted the A – B conditions to obtain the OXTR signal alone, and the A – C conditions to obtain the AVPR1a signal alone. We used this same approach for the AVPR1a radioligand (and the 3 associated conditions).

### 2.5 Statistical Analysis

#### 2.5.1 Hypothalamus

##### 2.5.1.1 Repeated Measures ANOVA for Radioligand Binding Specificity in the Hypothalamus

Two, one way RM-ANOVAs were used to determine binding specificity of both the OXTR and AVPR1a radioligands for the hypothalamus. Factors for the OXTR radioligand one-way repeated RM-ANOVA included the A, B, and C conditions detailed in the previous section. We did not use the subtraction approach for the hypothalamus RM-ANOVA, as the background values for the hypothalamus were consistent and reliable. An alpha level set to p < .05 was used to determine statistical significance.

##### 2.5.1.2 Independent Samples T-Test for Examining Group Differences in OXTR and AVPR1a Densities in the Hypothalamus

Due to the low anatomical specificity of our specimens, our sample size was underpowered to examine the combined effects of Group, Age, and Sex on OXTR and AVPR1a densities in the hypothalamus. We excluded specimens for which no appreciable levels of OXTR or AVPR1a signal were observed (likely due to the variation in anatomical location of the specimens provided), resulting in excluding 14 donors from the OXTR analysis and 16 donors from the AVPR1a analysis. Our resulting sample size was 14 (7 SZ, 7 control) for the OXTR analysis and 12 (7 SZ, 5 control) for the AVPR1a analysis(see Supplementary Tables 2 and 3 for descriptions of the included specimens). Therefore, we used independent samples t-tests to focus on examining differences in OXTR and AVPR1a mean densities between the SZ and control groups.Due to the difficulties assessing normality and homogeneity of variance due to our sample size, Welch’s t-tests were used. An alpha level set to p < .05 was used to determine statistical significance.

#### 2.5.2 Superior Temporal Sulcus

##### 2.5.2.1 RM-ANOVA for Radioligand Binding Specificity in the STS

A two-way RM-ANOVA was used to determine binding specificity of both the OXTR and AVPR1a radioligands for the STS. Factors included radioligand (^125^I-OVTA or ^125^I-LVA) and the receptor signal (OXTR or AVPR1a) resulting from the subtractions described above. This condition subtraction approach was only used for the RM-ANOVA for the STS. Tukey’s multiple comparisons were made across all four condition combinations. An alpha level set to *p* < .05 was used to determine statistical significance.

##### 2.5.2.2 Hierarchical Repeated Measures Multilevel Model

We tested the effects of independent variables diagnosis (SZ, unaffected), age, and sex on OXTR and AVPR1a receptor binding densities using two models, while controlling for subject-to-subject variability in receptor binding densities. For the OXTR, we examined OXTR binding from the OXTR radioligand (^125^I-OVTA), and for the AVPR1a, we examined AVPR1a binding from the AVPR1a radioligand (^125^I-LVA). Two levels were included in the model: Level two variables included the group, age, and sex of donors. The level one variable included repeated observations (three binding density values from three separate tissue slices per donor). We excluded 4 donors from the OXTR analysis and 5 donors from the AVPR1a analysis (see Supplementary Tables 4 and 5 for descriptions of the included specimens). For the OXTR analysis one donor was excluded due to a lack of OXTR signal, two donors were excluded due to tissue damage (freezing artifacts), and one donor was excluded due to film artifacts, resulting in a total of 37 donors. For the AVPR1a analysis, one donor was excluded due to tissue damage, and the remaining 4 donors were excluded due to film artifacts resulting in a total of 36 donors.

###### Model fit

A bottom-up approach was used to select the model with the best fit. We fit a series of models starting with a null model. The null model provided the intra-class correlation. In the model that included OXTR densities as the level one variable, the intra-class correlation indicated that 10% of variation in OXTR densities is due to donor-to-donor variability. In the model that included AVPR1a densities as the outcome variable, 86% of the variation in AVPR1a binding densities is due to donor-donor variability. Following the initial null model, we incorporated the predictors, adhering to the following sequence: 1) Integration of level 1 predictors into fixed maximum likelihood models. 2) Augmentation of fixed maximum likelihood models through the inclusion of level 2 predictors. 3) Specification of restricted maximum likelihood models, encompassing fixed effects. 4) Extension of the models to include both fixed and random effects within a restricted maximum likelihood framework. 5) Identification of the optimal model, distinguished by performance and utilizing restricted maximum likelihood. From the second step onward, models with the best fit were chosen before advancing to subsequent stages. The selection process was guided using log likelihood ratio tests (conducted using ANOVA version 4.0.2^25^), Akaike information criterion, Bayesian information criterion, and R-squared metrics and the systematic ranking of models based on their performance (performance version 0.5.0^26^). Each model included a random intercept to account for individual variability in receptor binding densities among donors. To evaluate the significance of pairwise differences, p-values were determined through t-tests using Satterthwaite degrees of freedom approximations. All analysis were conducted in R 4.2.1^27^.The *lme4* package (version 1.1-23^28^) as used to run hierarchical repeated measures models. A significance level of 0.05 was used throughout unless otherwise stated. All code and output are accessible at Open Science Framework (https://osf.io/7k4dv/).

## 3. Results

### 3.1 Superior Temporal Sulcus

#### 3.1.1 Competitive Binding Receptor Autoradiography

A two-way RM-ANOVA showed a significant main effect of radioligand *F*(1,142)= 16.12, *p*< .001) and a significant interaction between the radioligand and the receptor signal *F*(1,142) = 4.43, *p* = .037). Tukey’s multiple comparisons tests indicated significantly higher binding densities for the AVPR1a signal from ^125^I-LVA relative to all other conditions (Figure 2): compared to the OXTR signal from ^125^I-LVA (*p* = .0130), compared to the AVPR1a signal from ^125^I-OVTA (*p*< .001), and compared to the OXTR signal from ^125^I-OVTA (*p* = .002). The OXTR signal from ^125^I-LVA, the AVPR1a signal from ^125^I-OVTA, and the OXTR signal from ^125^I-OVTA were not significantly different from one another. Taken together, these results indicate that the predominant receptor subtype present in the STS is AVPR1a, with very minimal levels of OXTR.

**Figure 2.**
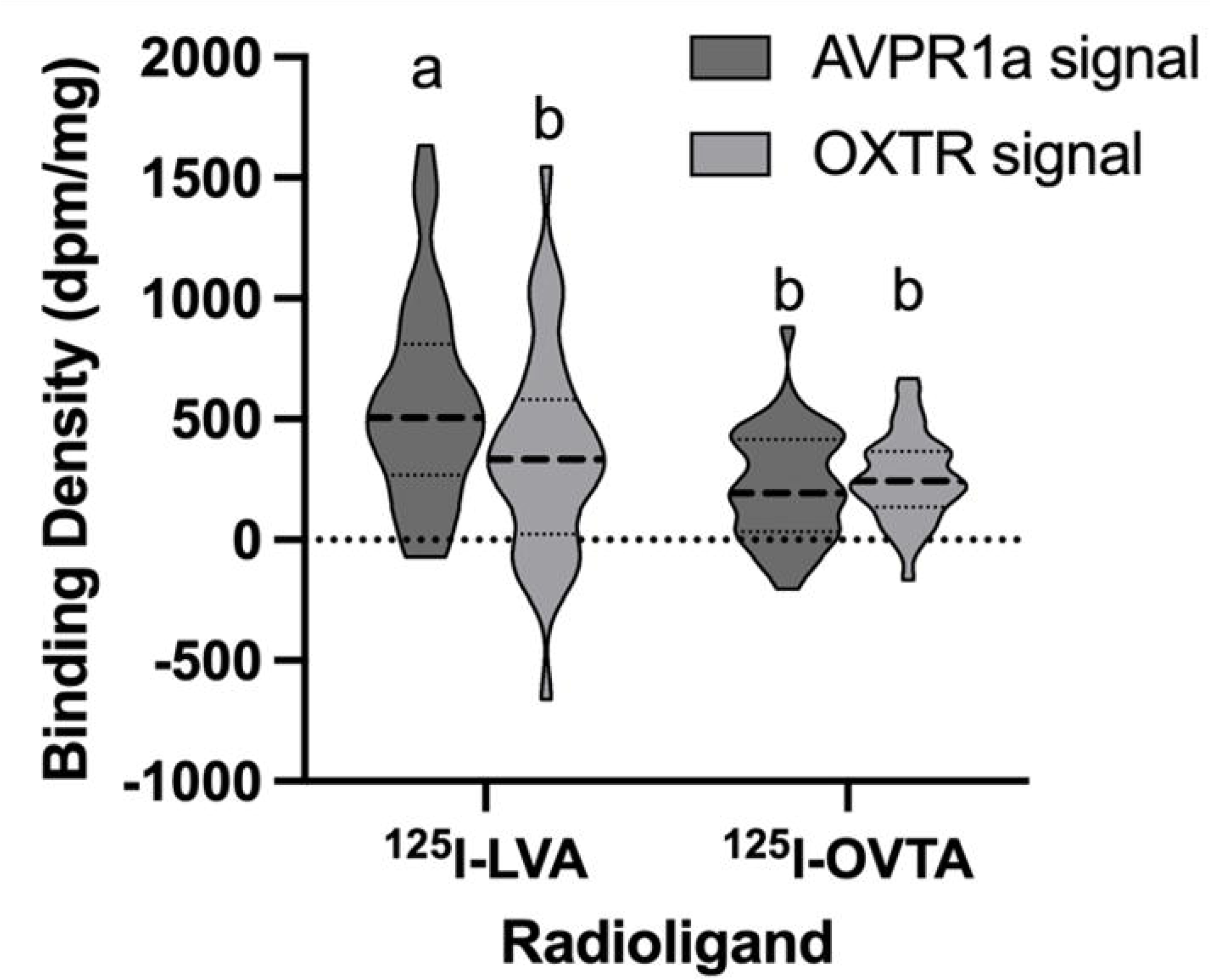
Receptor Binding Specificity in the Superior Temporal Sulcus Subjected to RM-ANOVA by Radioligands and Signal Type. The mean binding densities per condition here are after subtraction of the confounding signal. ^125^I-LVA = AVPR1a radioligand; ^125^I-OVTA = OXTR radioligand. The AVPR1a signal from ^125^I-LVA is significantly different (a) from all other conditions, which do not significantly differ from each other (b).

#### 3.1.2 AVPR1a Receptor Density

The final hierarchical repeated measures model showed a significant three-way interaction between Group, Age, and Sex on AVPR1a receptor density in STS, *t*(28.03) = −2.74, *p* = .010 (Table 1).

**Table 1.**
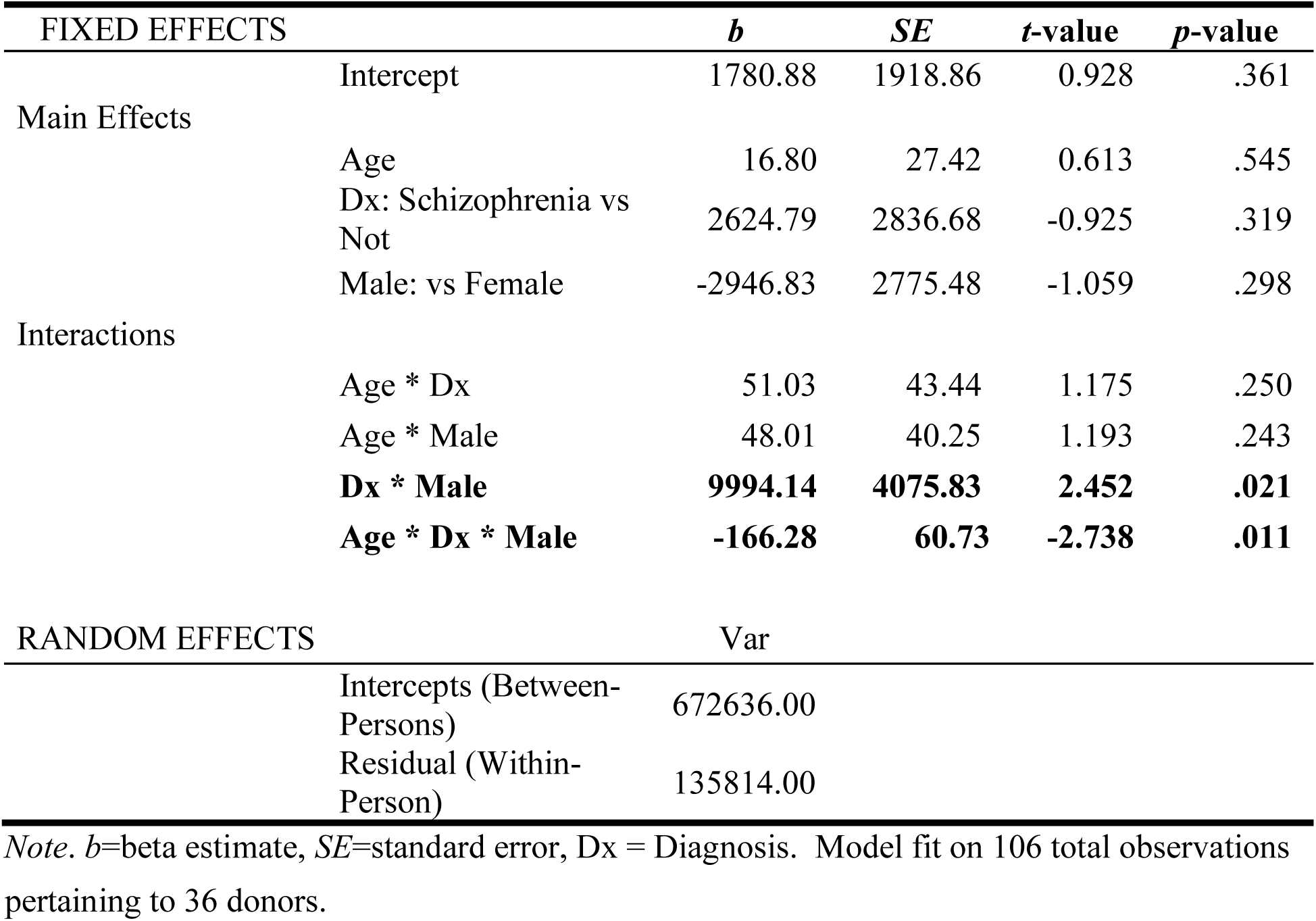
Parameter Estimates for Hierarchical Repeated Measures Multilevel Regression Model of Vasopressin Receptor Densities on Diagnosis of Schizophrenia, Moderated by Both Age and Sex.

The combined effect of Group and Age on AVPR1a receptor density depends on Donor Sex (Figure 3). In the unaffected control group, AVPR1a receptor binding increases with age in males but not females. In the SZ group, females show an increase in AVPR1a receptor binding with age, whereas males show the opposite effect. Simple slopes analysis indicated a 1-year increase in age in male controls is associated with a 64.81 (dpm/mg) increase in AVPR1a receptor binding, *SE* = 29.38, *p* = .036. In females who had SZ, a 1-year increase in age is associated with a 67.83 (dpm/mg) increase in AVPR1a receptor binding, *SE*= 33.69, *p* = .054. Although the p-value is right at the threshold of reaching statistical significance, we interpret this as evidence to support a biologically meaningful change in AVPR1a in the female SZ group. However, our findings should be interpreted with caution, as one female donor in the SZ group with very high AVPR1a receptor binding (Figure 3) appears to be driving this effect. When removing this donor from the analysis, the three-way interaction is no longer present. Upon further examination of the donor’s records (Supplementary Table 6) and AVPR1a binding in the autoradiogram (Supplementary Figure 1), we did not find any reasonable cause to exclude this donor from our analysis. It is possible that the increase in AVPR1a receptor density with age in SZ females is a true effect, however, our sample is limited due to the lack of female donors who had SZ within the ages of 66 to 83.

**Figure 3.**
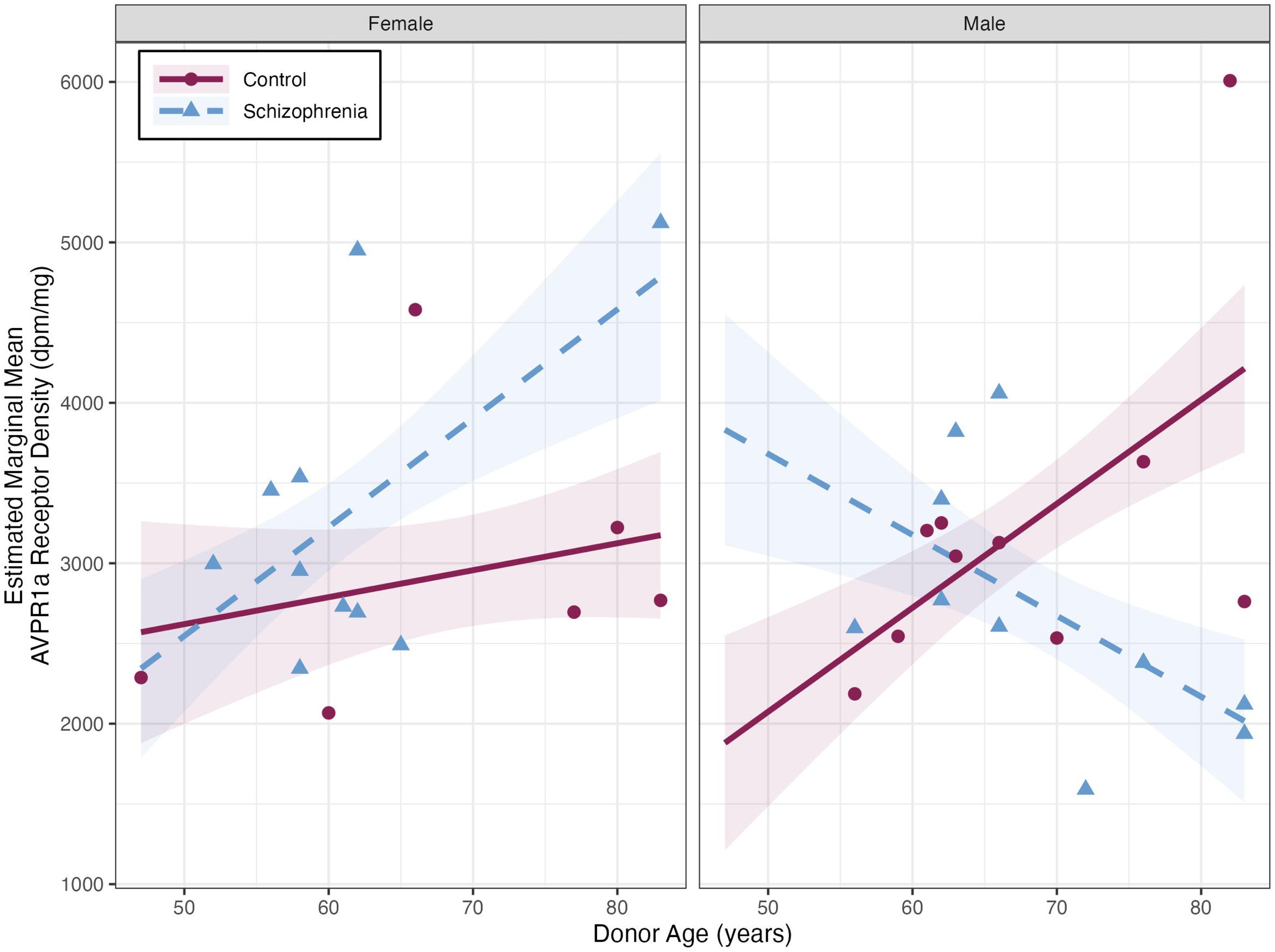
Combined Effect of Diagnosis, Sex, and Age on AVPR1a Receptor Densities in the Superior Temporal Sulcus. Estimated marginal mean AVPR1a receptor binding density (dpm/mg) based on donor group, sex, and age. Bands represent +/− 1 standard error of the mean.

In the subset of specimens for which antipsychotic medication history was available (n = 8), we found no significant correlation between STS AVPR1a densities and antipsychotic dosages, suggesting our findings were not due to medication effects (Supplementary Figure 2).

#### 3.1.3 OXTR Receptor Density

The best fitting model determined by the likelihood ratio test, showed a significant main effect of Age, *t*(32.23)= 2.23, *p* = .033, χ^2^ (6) = 5.16, *p* = .023. When controlling for Donor Group, Sex, and donor-to-donor variability in receptor binding, each 1-year increase in age is associated with a 6.69 (dpm/mg; +/− 3 standard error) increase in OXTR binding density (Figure 4).We again found no significant correlation between STS OXTR densities and antipsychotic dosages (n = 8), suggesting a lack of medication effects (Supplementary Figure 3).

**Figure 4.**
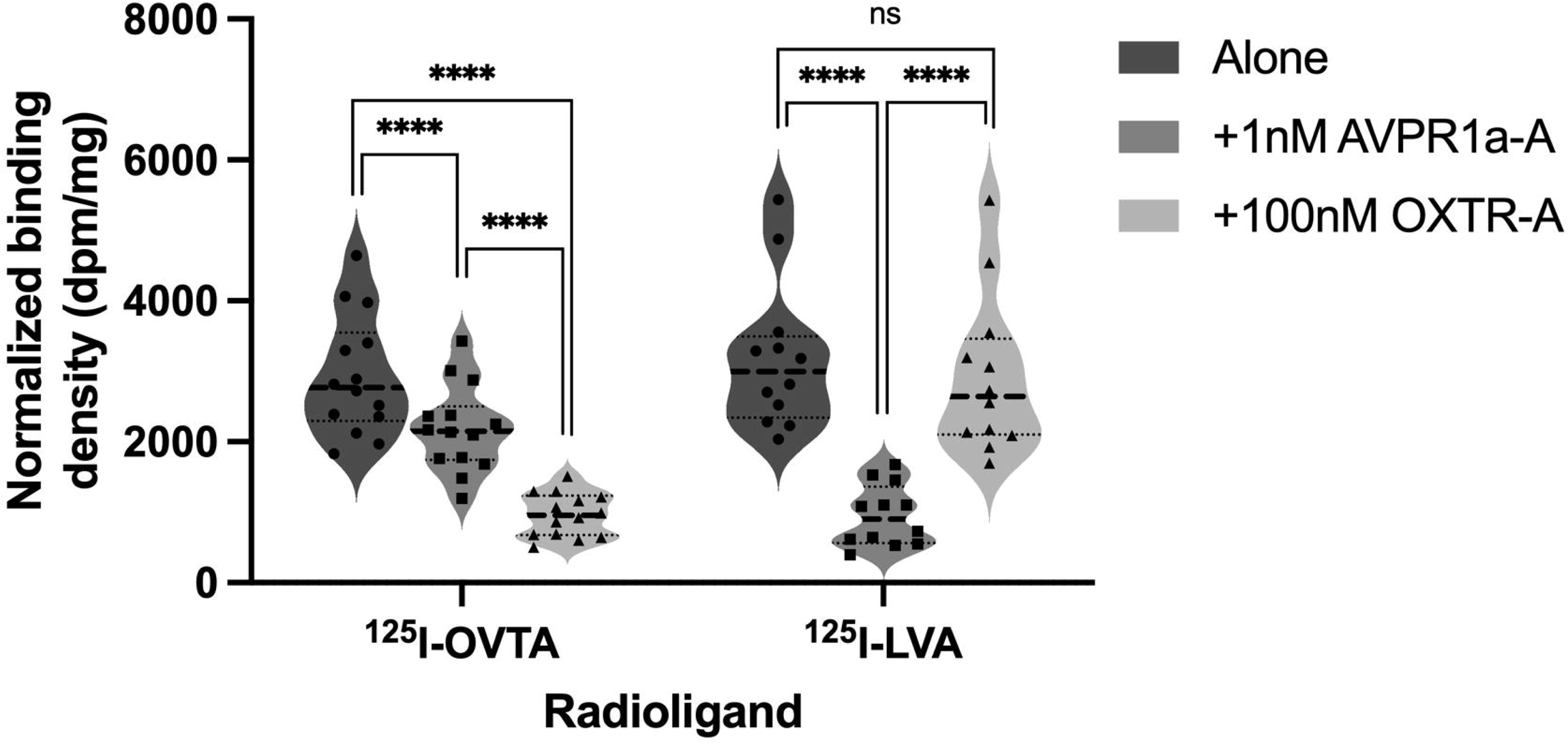
Competitive Binding Results for each Radioligand in the Hypothalamus. A) Binding of the OXTR radioligand in the hypothalamus in all three competition conditions. B) Binding of the AVPR1a radioligand in the hypothalamus in all three competition conditions. Dashed lines indicate mean.Dotted lines represented +/− 1 standard error of the mean. **** = p < .001.

### 3.2 Hypothalamus

#### 3.2.1 Competitive Binding Receptor Autoradiography

A one-way repeated measures ANOVA showed a significant effect of Condition on binding densities, *F*(1.28, 16.70)=62.44, *p*< . 001 (Greenhouse-Geisser correction for violated assumption of sphericity), between OXTR alone (*M*= 2921) compared to +1nM AVPR1a-A (*M*= 2186) and +100nm OXTR-A (*M*= 1967; Figure 4A). A one-way repeated measures ANOVA showed a significant effect of Condition on binding densities, *F*(1.31, 14.40)=47.58, *p*< . 001 (Greenhouse-Geisser correction for violated assumption of sphericity), between AVPR1a alone (*M*=3182) compared to +1nM AVPR1a-A (*M*= 951.2) and +100nm OXTR-A (*M*= 2918; Figure 4B). Representative autoradiograms from this competitive binding approach are provided in the supplementary materials (Supplementary Figures 4 and 5).

#### 3.2.2 Effects of Group on OXTR and AVPR1a Densities

An independent samples t-test revealed a significant difference in mean OXTR densities in the hypothalamus between the SZ and unaffected control groups (Figure 5A), *t*(11.17) = −2.26, *p* = .044. Higher OXTR densities were observed in SZ (*M* = 2507.97) versus unaffected controls (*M* =1863.95).The independent samples t-test revealed a trend toward elevated AVPR1a densities in SZ (*M* = 3373.35) relative to unaffected controls (*M* = 2280.39), *t*(7.14) = −3.69, *p* = .069 (Figure 5B).Due to the low number of donors in the hypothalamus analysis for which medication records were available(OXTR: n = 3 and AVPR1a: n=4), we were unable to assess correlations between antipsychotic medication dosage and OXTR or AVPR1a densities.

**Figure 5.**
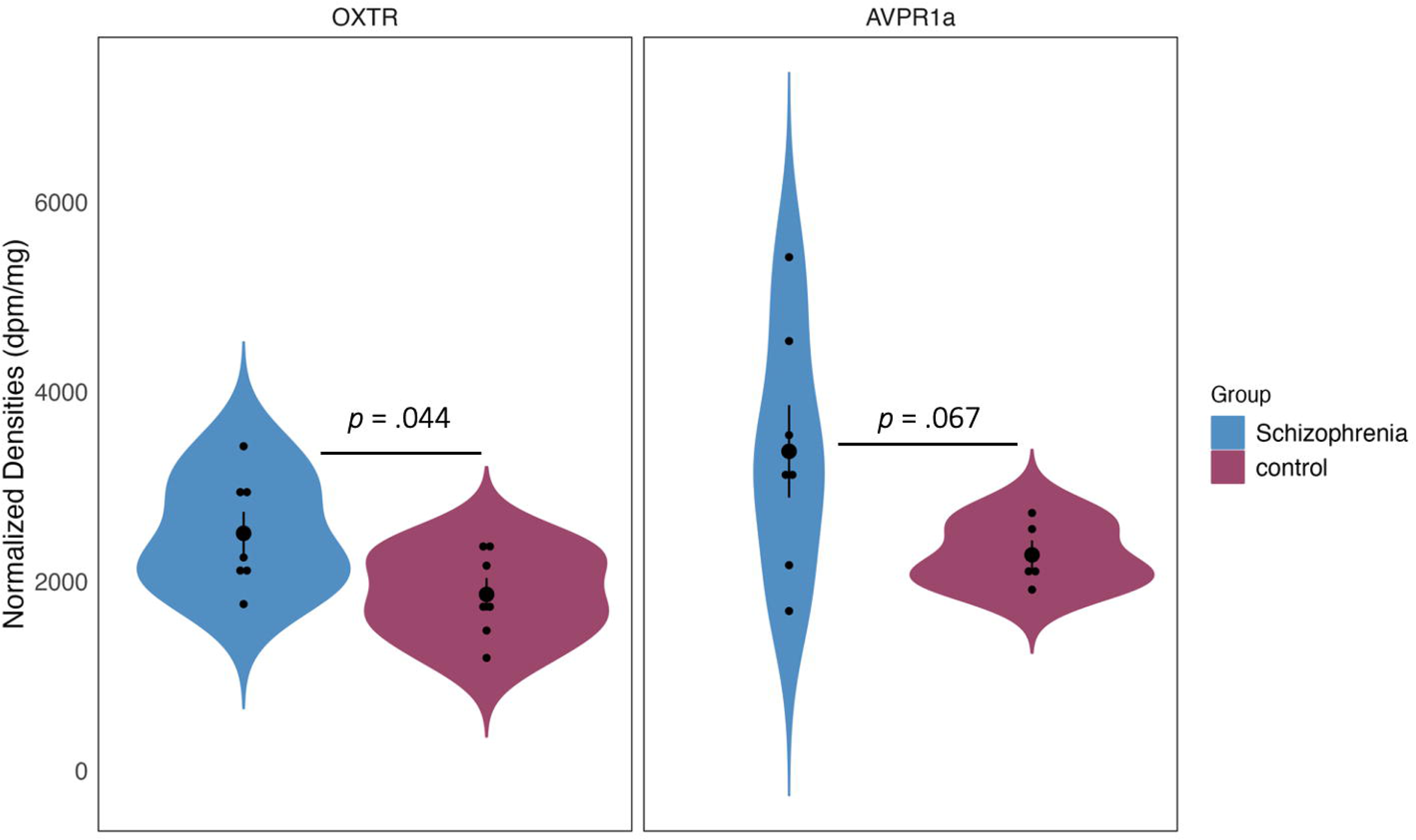
Elevated OXTR (A)and AVPR1a (B) Densities in Schizophrenia in the Hypothalamus. Means per each diagnostic group after baseline subtraction. Vertical lines represent +/− 1 standard error of the mean.

## 4. Discussion

Using competitive binding receptor autoradiography to selectively quantify OXTR and AVPR1a, we found increased OXTR densities and evidence of increased AVPR1a densities in the hypothalamus in SZ. In the STS of females who had SZ, AVPR1a density increases with age. Unaffected male controls also showed increased AVPR1a density with age in the STS. It is possible that our finding in SZ females was driven by one 83-year-old donor, so future postmortem studies should include larger samples between the ages of 65 to 80. In the current study, female donors within this age bracket did not exist in the brain bank with our inclusion and exclusion criteria.

### 4.1 Effect of Schizophrenia on OXTR and AVPR1a Densities in the Hypothalamus

Our finding of increased hypothalamic OXTR in SZ aligns with our hypothesis and underscores the potential role of OXT in regulating the HPA axis. Ventromedial hypothalamic OXT releaseexerts anxiolytic effects during negative social interactions, such as social defeat^3^, where rodents exhibit submissive behavior following exposure to aggressive conspecifics. Furthermore, intracerebroventricular OXT infusion in rodents mitigates plasma corticosterone release in response to noise stress^29^, suggesting OXTergic modulation of HPA activity during stressful events. Our observation of elevated hypothalamic OXTR densities in SZ individuals may compensate for reduced endogenous release of hypothalamic OXT. This potential deficiency in OXT release could disinhibit HPA activity and lead to upregulated hypothalamic OXTR as well as HPA hyperactivity. HPA hyperactivity is observed in SZ, as evidenced by elevated baseline cortisol levels^23^and enlarged hypothalamic volumes^21^. In the context of limited endogenous OXT release, particularly during socially and psychologically stressful events, the capacity for OXT to serve as a social buffer may be compromised in SZ. Consequently, individuals with SZ may perceive negative, or particularly stress-inducing social interactions with heightened salience.

We also observed a trend (*p* = .069) towards increased AVPR1a densities in the hypothalamus in SZ relative to controls. This trend does not support our hypothesis and contrasts with previous findings reporting reduced *AVPR1a* mRNA expression in SZ^22^. In a previous study, the paraventricular nucleus showed diminished AVPR1a mRNA in SZ^22^. However, altered *AVPR1a* mRNA was specific to the paraventricular nucleus, with no observed differences in the supraoptic nucleus. Our study lacked such anatomical specificity due to limitations in specimen collection, preventing us from analyzing specific hypothalamic subregions. A previous study reported a diminished peripheral AVP/adrenocorticotropin stress response in SZ^30^. Sustained blunting of peripheral AVP in response to stress could contribute to upregulated AVPR1a in the hypothalamus over time. In future studies, it will be valuable to investigate both OXT and AVP densities and mRNA expression within specific hypothalamic subregions to directly assess receptor gene expression and protein levels.

### 4.2 Selective binding of AVPR1a in the superior temporal gyrus

To our knowledge, this is the first study to report the presence of AVPR1a receptors in the temporal cortex in the human brain. This finding is contrary to early work reporting no specific binding of OXTR or AVPR1a in the frontal and temporal cortices^14^. Due to the affinity of these radioligands for both OXTR and AVPR1a, the binding reported by Loup and colleagues^14^ likely includes signal from both receptors. In addition, the ^3^H-AVP radioligand previously used in that study has low selectivity to AVPR1a relative to the iodinated ligand used in the current study (^125^I-LVA), which could have resulted in binding to OXTR in some regions. Our findings are, however, consistent with primate and human studies using ^125^I-LVA and ^125^I-OVTA, which report minimal OXTR binding in the cortex, yet abundant, widely dispersed AVPR1a binding in cortical areas^31^.

The STS is highly implicated in social perception and appears to be recruited by IN-OXT during theory of mind tasks in SZ^18^. Our findings suggest that STS recruitment by IN-OXT and the associated social cognitive effects in SZ likely occurs by binding of exogenous OXT to AVPR1a receptors in the STS rather than the OXTR. Mechanistic action of exogenous OXT binding to AVPR1a (and vice versa) is supported by animal studies^32^. For example, in OXTR knockout mice, intracerebroventricular infusion of OXT or AVP both restore social exploration and social recognition^32^. It is important to note that our interpretation of the effects of IN-OXT on AVPR1a in the STS assumes that administration of IN-OXT or IN-AVP crosses the blood brain barrier, which has yet to be decisively shown in humans. However, a recent primate study confirmed that IN-OXT can be detected in several brain regions including the striatum, cerebellum, thalamus, and cortical areas^33^.

### 4.3 Effect of Diagnosis, Sex, and Age on AVPR1a receptor binding

We found an effect of diagnosis, sex, and age on AVPR1a densities. AVP is synthesized in the paraventricular and supraoptic nuclei of the hypothalamus^34^, and is integrated in the HPA axis. AVP released from parvocellular neurons in the paraventricular nucleus stimulates secretion of adrenocorticotropic hormone from the anterior pituitary and cortisol from the adrenal glands^34^.This coordinated system facilitates social behavior when faced with a stressor, including aggression (fight) and avoidance (flight).

We found increased AVPR1a density with advanced age in women who had SZ.Due to AVP’s well established role in modulating social stress^34^ and aggression^5^in animals, increased AVPR1a binding in the STS with age could explain sex differences in symptoms observed in older adults with SZ. Gur and colleagues^35^ found that sex differences in symptom severity were age dependent. In people with SZ between the ages of 65 to 80, more severe negative symptoms were observed in men relative to women. However, in women over the age of 80, negative symptoms and suspiciousness/hostility scores were worse relative to men. Therefore, upregulated AVPR1a with advanced age in females with SZ may contribute to enhanced suspicion toward others and more negative symptoms (e.g., social withdrawal). Elevated AVPR1a densities in women with SZ may reflect a threshold for detecting threat in social information, a task for which the STS is fine-tuned. Unfortunately, clinical symptom scores were only available for two donors, precluding assessment of relationships between symptom severity and receptor binding densities due tolow sample size. Future studies with clinical records are needed to determine the relationship between AVPR1a densities and symptom severity.

Sex dependent effects on AVPR1a in SZ could also be related to changes in estrogen with advanced age. Estrogen is believed to have protective effects in women with SZ; the greatest vulnerability for onset of SZ in women is during periods across the lifespan when estrogen is low (e.g., menopause^36^). Importantly, this peak in illness onset during this stage of life (e.g., ages 45 to 55) is not observed in men^36^. In addition, women with SZ show a greater risk of hospitalization for psychotic symptoms during phases of the menstrual cycle when estrogen is low^36^. Lower peripheral estradiol concentrations are observed in women with SZ relative to controls^37^. Estrogen modulates AVP and OXT neuronal activity via the estrogen receptorβ^38^. Older adults receiving continuous estrogen treatment show upregulated *AVPR1a* expression^39^. However, estrogen’s effects on *AVPR1a* gene expression are region dependent, as estrogen treatment may also decrease *AVPR1a* expression in the arcuate nucleus of rats^40^. Given estrogen’s ability to up or downregulate *AVPR1a* expression, an accelerated decline in estrogen with age in women with SZ may contribute to upregulated AVPR1a densities. However, the effects of advanced age on estrogen levels in the brain are unknown in SZ. Future studies co-localizing *AVPR1a* and *estrogen receptor β* mRNA are needed to determine the relationship between these two systems in the SZ brain.

### 4.4 Effect of age on AVPR1a binding in male controls

Our study found that male controls showed increased AVPR1a densities in the STS with advanced age. A previous study indicated that IN-AVP increased right STS activation during the formation of competitive decisions in response to provocation^41^, suggesting enhanced mentalizing before social interactions. IN-AVP affects social behavior and neural activation in a sex-dependent manner. In men, but not women, IN-AVP increases cooperation and activates regions including the nucleus accumbens and amygdala^42^. Additionally, IN-AVP reduces the STS/temporal parietal junction response to negative social feedback in males^43^, indicating its role in enhancing social perception or buffering negative emotions. Therefore, AVP either recruits brain regions necessary for social perception or diminishes neural activation to buffer the emotionally aversive experience of negative social interactions. Our findings suggest that increased STS AVPR1a in older men may enhance the STS response to IN-AVP during social interactions. Further research is needed to confirm age and sex-dependent effects of IN-AVP on neural activation and social behavior in older adults.

Rodent studies highlight the interplay between androgens and AVPR1a expression. In male hamsters, adult gonadectomy decreased AVPR1a densities in many social regions including the bed nucleus of the stria terminalis and the ventromedial hypothalamus^44^. These brain regions are implicated in aggressive and territorial behaviors in rodents, such as flank marking^44^. In Syrian hamsters, treatment with anabolic steroids during adolescence increases AVPR1a binding in key social brain regions, including the lateral septum, ventrolateral hypothalamus, and bed nucleus of the stria terminalis^45^. While DeLeon and colleagues (2002) specifically investigate the effects of anabolic steroid exposure during adolescence in hamsters, their study provides evidence that androgens can influence AVPR1a binding within the brain. Given that testosterone levels decline with age in men, the findings from animal work suggest a possible mechanism by which age-related hormonal changes influence a compensatory mechanism resulting in upregulated AVPR1a receptor expression with advanced age in male humans, as reported here. However, further research is needed to directly investigate the implications of sex and age-related hormonal changes on AVPR1a receptor expression in the human brain.

### 4.5 OXTR Binding

No significant differences in OXTR densities in the STS between SZ and controls were observed. This finding is consistent with previous research reporting no differences in OXTR binding in the temporal cortex between the SZ and control groups^15^. It is important to note that Uhrig and colleagues^15^ analyzed Brodmann’s area 21, the medial temporal gyrus which lies adjacent and ventral to the superior temporal gyrus. Due to the methodological limitations of using the ^125^I-OVTA radioligand to visualize OXTR in human brain tissue^13^, the OXTR binding analyzed by Uhrig and colleagueslikely includes confounding AVPR1a binding. In any case, our findings corroborate the lack of differences in mean OXTR binding between SZ and controls even when removing confounding AVPR1a signal.

Interestingly, although no effects of diagnosis on OXTR binding was observed in the medial temporal gyrus, Uhrig and colleagues^15^ found reduced *OXTR* mRNA in SZ v. controls. This discrepancy between OXTR binding and OXTR mRNA levels may be due to the ability of mRNA to be transported to axons after the gene is transcribed, which has been observed in *OXTR* mRNA in the rodent hypothalamus^46^. Future studies examining both OXTR binding densities and mRNA are needed to determine variation in the OXTR at both the protein and genetic levels.

Although we found no significant effects of diagnosis on OXTR binding, we observed an increase in OXTR binding densities with age across all donors. This statistically significant finding may be less biologically meaningful due to the low levels of OXTR binding detected in the STS. However, this upregulation aligns with reports of increased OXTR gene expression in the aging human brain^47^. According to Socioemotional Selectivity Theory, aging leads to more selective social decisions, resulting in smaller social networks^48^. Older adults also exhibit a positivity bias and increased trust^49^. Upregulated OXTR in the STS may contribute to these shifts in sociality with age and heightened sensitive to trust during face evaluation.

## 5. Conclusions

We found increased OXTR and evidence of increased AVPR1a densities in the hypothalamus in SZ relative to controls. Upregulated OXTR and AVPR1a may be compensation for low endogenous hypothalamic release of these neurohormones in SZ. SZ females showed increased STS AVPR1a binding with advanced age. Upregulated AVPR1a with advanced age may drive increases in negative symptom severity observed in women with SZ.

## Declaration of Interest

The authors have no conflicts of interest to report.

## Acknowledgements

Funding was provided by the Office of Research at Utah State University. The authors would like to thank Gentry Mower, Logan Hale, and Mckenna Rich for specimen preparation assistance.

## Notes

https://osf.io/7k4dv

### Competing Interest Statement

The authors have declared no competing interest.

## References

1. McGrath, J., Saha, S., Chant, D., & Welham, J. (2008). Schizophrenia: A Concise Overview of Incidence, Prevalence, and Mortality. Epidemiologic Reviews, 30(1), 67–76. 10.1093/epirev/mxn001

2. Green, M. F. (2016). Impact of cognitive and social cognitive impairment on functional outcomes in patients with schizophrenia. The Journal of Clinical Psychiatry, 77 Suppl 2, 8–11. 10.4088/JCP.14074su1c.02

3. Veenema, A. H., & Neumann, I. D. (2008). Central vasopressin and oxytocin release: Regulation of complex social behaviours. Progress in Brain Research, 170, 261–276. 10.1016/S0079-6123(08)00422-6

4. Meyer-Lindenberg, A., Domes, G., Kirsch, P., & Heinrichs, M. (2011). Oxytocin and vasopressin in the human brain: Social neuropeptides for translational medicine. Nature Reviews Neuroscience, 12(9), 524–538. 10.1038/nrn3044

5. Albers, H. E. (2012). The regulation of social recognition, social communication and aggression: Vasopressin in the social behavior neural network. Hormones and Behavior, 61(3), 283–292. 10.1016/j.yhbeh.2011.10.007

6. Heinrichs, M., von Dawans, B., & Domes, G. (2009). Oxytocin, vasopressin, and human social behavior. Frontiers in Neuroendocrinology, 30(4), 548–557. 10.1016/j.yfrne.2009.05.005

7. Shilling, P. D., & Feifel, D. (2016). Potential of Oxytocin in the Treatment of Schizophrenia. CNS Drugs, 30(3), 193–208. 10.1007/s40263-016-0315-x

8. Rubin, L. H., Carter, C. S., Bishop, J. R., Pournajafi-Nazarloo, H., Harris, M. S. H., Hill, S. K., Reilly, J. L., & Sweeney, J. A. (2013). Peripheral vasopressin but not oxytocin relates to severity of acute psychosis in women with acutely-ill untreated first-episode psychosis. Schizophrenia Research, 146(1), 138–143. 10.1016/j.schres.2013.01.019

9. Rubin, L. H., Carter, C. S., Bishop, J. R., Pournajafi-Nazarloo, H., Drogos, L. L., Hill, S. K., Ruocco, A. C., Keedy, S. K., Reilly, J. L., Keshavan, M. S., Pearlson, G. D., Tamminga, C. A., Gershon, E. S., & Sweeney, J. A. (2014). Reduced Levels of Vasopressin and Reduced Behavioral Modulation of Oxytocin in Psychotic Disorders. Schizophrenia Bulletin, 40(6), 1374–1384. 10.1093/schbul/sbu027

10. Aydın, O., Lysaker, P. H., Balıkçı, K., Ünal-Aydın, P., & Esen-Danacı, A. (2018). Associations of oxytocin and vasopressin plasma levels with neurocognitive, social cognitive and meta cognitive function in schizophrenia. Psychiatry Research, 270, 1010–1016. 10.1016/j.psychres.2018.03.048

11. Leng, G., & Sabatier, N. (2016). Measuring Oxytocin and Vasopressin: Bioassays, Immunoassays and Random Numbers. Journal of Neuroendocrinology, 28(10), jne.12413. 10.1111/jne.12413

12. Freeman, S. M., Samineni, S., Allen, P. C., Stockinger, D., Bales, K. L., Hwa, G. G. C., & Roberts, J. A. (2016). Plasma and CSF oxytocin levels after intranasal and intravenous oxytocin in awake macaques. Psychoneuroendocrinology, 66, 185–194. 10.1016/j.psyneuen.2016.01.014

13. Freeman, S. M. (2022). Using Receptor AutoradiographyAutoradiography to Visualize and Quantify OxytocinOxytocinAutoradiographyVasopressinsAutoradiographyand VasopressinVasopressins1a Receptors in the Human and Nonhuman PrimatePrimatesBrain. In E. L. Werry, T. A. Reekie, & M. Kassiou (Eds.), Oxytocin: Methods and Protocols (pp. 105–125). Springer US. 10.1007/978-1-0716-1759-5_7

14. Loup, F., Tribollet, E., Dubois-Dauphin, M., & Dreifuss, J. J. (1991). Localization of high-affinity binding sites for oxytocin and vasopressin in the human brain. An autoradiographic study. Brain Research, 555(2), 220–232. 10.1016/0006-8993(91)90345-V

15. Uhrig, S., Hirth, N., Broccoli, L., von Wilmsdorff, M., Bauer, M., Sommer, C., Zink, M., Steiner, J., Frodl, T., Malchow, B., Falkai, P., Spanagel, R., Hansson, A. C., & Schmitt, A. (2016). Reduced oxytocin receptor gene expression and binding sites in different brain regions in schizophrenia: A post-mortem study. Schizophrenia Research, 177(1–3), 59–66. 10.1016/j.schres.2016.04.019

16. Hein, G., & Knight, R. T. (2008). Superior Temporal Sulcus—It’s My Area: Or Is It? Journal of Cognitive Neuroscience, 20(12), 2125–2136. 10.1162/jocn.2008.20148

17. Yan, Z., Schmidt, S. N. L., Frank, J., Witt, S. H., Hass, J., Kirsch, P., & Mier, D. (2020). Hyperfunctioning of the right posterior superior temporal sulcus in response to neutral facial expressions presents an endophenotype of schizophrenia. Neuropsychopharmacology, 45(8), 1346–1352. 10.1038/s41386-020-0637-8

18. De Coster, L., Lin, L., Mathalon, D. H., & Woolley, J. D. (2019). Neural and behavioral effects of oxytocin administration during theory of mind in schizophrenia and controls: A randomized control trial. Neuropsychopharmacology, 44(11), 1925–1931. 10.1038/s41386-019-0417-5

19. Abram, S. V., De Coster, L., Roach, B. J., Mueller, B. A., van Erp, T. G. M., Calhoun, V. D., Preda, A., Lim, K. O., Turner, J. A., Ford, J. M., Mathalon, D. H., & Woolley, J. D. (2020). Oxytocin Enhances an Amygdala Circuit Associated With Negative Symptoms in Schizophrenia: A Single-Dose, Placebo-Controlled, Crossover, Randomized Control Trial. Schizophrenia Bulletin, 46(3), 661–669. 10.1093/schbul/sbz091

20. Saper, C. B., & Lowell, B. B. (2014). The hypothalamus. Current Biology, 24(23), R1111–R1116. 10.1016/j.cub.2014.10.023

21. Tognin, S., Rambaldelli, G., Perlini, C., Bellani, M., Marinelli, V., Zoccatelli, G., Alessandrini, F., Pizzini, F. B., Beltramello, A., Terlevic, R., Tansella, M., Balestrieri, M., & Brambilla, P. (2012). Enlarged hypothalamic volumes in schizophrenia. Psychiatry Research: Neuroimaging, 204(2), 75–81. 10.1016/j.pscychresns.2012.10.006

22. Busch, J. R., Jacobsen, C., Lynnerup, N., Banner, J., & Møller, M. (2019). Expression of vasopressin mRNA in the hypothalamus of individuals with a diagnosis of schizophrenia. Brain and Behavior, 9(9). 10.1002/brb3.1355

23. Ritsner, M., Gibel, A., Maayan, R., Ratner, Y., Ram, E., Modai, I., & Weizman, A. (2007). State and trait related predictors of serum cortisol to DHEA(S) molar ratios and hormone concentrations in schizophrenia patients. European Neuropsychopharmacology, 17(4), 257–264. 10.1016/j.euroneuro.2006.09.001

24. Windle, R. J. (2004). Oxytocin Attenuates Stress-Induced c-fos mRNA Expression in Specific Forebrain Regions Associated with Modulation of Hypothalamo-Pituitary-Adrenal Activity. Journal of Neuroscience, 24(12), 2974–2982. 10.1523/JNEUROSCI.3432-03.2004

25. Cleveland, W. S., Grosse, E., Shyu, W. M., Chambers, J. M., & Hastie, T. J. (1992). Statistical models in S. Local Regression Models, Chapter-8.

26. Nakagawa, S., Johnson, P. C. D., & Schielzeth, H. (2017). The coefficient of determination *R*^2^ and intra-class correlation coefficient from generalized linear mixed-effects models revisited and expanded. Journal of The Royal Society Interface, 14(134), 20170213. 10.1098/rsif.2017.0213

27. R Core Team. (2023). R: A Language and Environment for Statistical Computing. R Foundation for Statistical Computing. https://www.R-project.org/

28. Bates, D., Maechler, M., Bolker, B., Walker, S., Christensen, R. H. B., Singmann, H., Dai, B., Grothendieck, G., Green, P., & Bolker, M. B. (2015). Package ‘lme4.’ Convergence, 12(1), 2.

29. Windle, R. J., Shanks, N., Lightman, S. L., & Ingram, C. D. (1997). Central Oxytocin Administration Reduces Stress-Induced Corticosterone Release and Anxiety Behavior in Rats*. Endocrinology, 138(7), 2829–2834. 10.1210/endo.138.7.5255

30. Goldman, M. B., Gnerlich, J., & Hussain, N. (2007). Neuroendocrine Responses to a Cold Pressor Stimulus in Polydipsic Hyponatremic and in Matched Schizophrenic Patients. Neuropsychopharmacology, 32(7), 1611–1621. 10.1038/sj.npp.1301282

31. Freeman, S. M., & Young, L. J. (2016). Comparative Perspectives on Oxytocin and Vasopressin Receptor Research in Rodents and Primates: Translational Implications. Journal of Neuroendocrinology, 28(4). 10.1111/jne.12382

32. Sala, M., Braida, D., Lentini, D., Busnelli, M., Bulgheroni, E., Capurro, V., Finardi, A., Donzelli, A., Pattini, L., Rubino, T., Parolaro, D., Nishimori, K., Parenti, M., & Chini, B. (2011). Pharmacologic Rescue of Impaired Cognitive Flexibility, Social Deficits, Increased Aggression, and Seizure Susceptibility in Oxytocin Receptor Null Mice: A Neurobehavioral Model of Autism. Biological Psychiatry, 69(9), 875–882. 10.1016/j.biopsych.2010.12.022

33. Lee, M. R., Shnitko, T. A., Blue, S. W., Kaucher, A. V., Winchell, A. J., Erikson, D. W., Grant, K. A., & Leggio, L. (2020). Labeled oxytocin administered via the intranasal route reaches the brain in rhesus macaques. Nature Communications, 11(1), 2783. 10.1038/s41467-020-15942-1

34. Landgraf, R., & Neumann, I. D. (2004). Vasopressin and oxytocin release within the brain: A dynamic concept of multiple and variable modes of neuropeptide communication. Frontiers in Neuroendocrinology, 25(3), 150–176. 10.1016/j.yfrne.2004.05.001

35. Gur, R. E., Petty, R. G., Turetsky, B. I., & Gur, R. C. (1996). Schizophrenia throughout life: Sex differences in severity and profile of symptoms. Schizophrenia Research, 21(1), 1–12. 10.1016/0920-9964(96)00023-0

36. Seeman, M. V. (2012). Women and Psychosis. Women’s Health, 8(2), 215–224. 10.2217/WHE.11.97

37. Markham, J. A. (2012). Sex steroids and schizophrenia. Reviews in Endocrine and Metabolic Disorders, 13(3), 187–207. 10.1007/s11154-011-9184-2

38. Winslow, J. T., & Insel, T. R. (2004). Neuroendocrine basis of social recognition. Current Opinion in Neurobiology, 14(2), 248–253. 10.1016/j.conb.2004.03.009

39. Garcia, A. N., Depena, C. K., Yin, W., & Gore, A. C. (2016). Testing the critical window of estradiol replacement on gene expression of vasopressin, oxytocin, and their receptors, in the hypothalamus of aging female rats. Molecular and Cellular Endocrinology, 419, 102–112. 10.1016/j.mce.2015.10.004

40. Yin, W., Maguire, S. M., Pham, B., Garcia, A. N., Dang, N.-V., Liang, J., Wolfe, A., Hofmann, H. A., & Gore, A. C. (2015). Testing the Critical Window Hypothesis of Timing and Duration of Estradiol Treatment on Hypothalamic Gene Networks in Reproductively Mature and Aging Female Rats. Endocrinology, 156(8), 2918–2933. 10.1210/en.2015-1032

41. Brunnlieb, C., Münte, T. F., Krämer, U., Tempelmann, C., & Heldmann, M. (2013). Vasopressin modulates neural responses during human reactive aggression. Social Neuroscience, 8(2), 148–164. 10.1080/17470919.2013.763654

42. Rilling, J. K., DeMarco, A. C., Hackett, P. D., Chen, X., Gautam, P., Stair, S., Haroon, E., Thompson, R., Ditzen, B., Patel, R., & Pagnoni, G. (2014). Sex differences in the neural and behavioral response to intranasal oxytocin and vasopressin during human social interaction. Psychoneuroendocrinology, 39, 237–248. 10.1016/j.psyneuen.2013.09.022

43. Gozzi, M., Dashow, E. M., Thurm, A., Swedo, S. E., & Zink, C. F. (2017). Effects of Oxytocin and Vasopressin on Preferential Brain Responses to Negative Social Feedback. Neuropsychopharmacology: Official Publication of the American College of Neuropsychopharmacology, 42(7), 1409–1419. 10.1038/npp.2016.248

44. Young, L. J., Wang, Z., Cooper, T. T., & Albers, H. E. (2000). Vasopressin (V1a) Receptor Binding, mRNA Expression and Transcriptional Regulation by Androgen in the Syrian Hamster Brain. Journal of Neuroendocrinology, 12(12), 1179–1185. 10.1046/j.1365-2826.2000.00573.x

45. DeLeon, K. R., Grimes, J. M., & Melloni, R. H. (2002). Repeated Anabolic-Androgenic Steroid Treatment during Adolescence Increases Vasopressin V1A Receptor Binding in Syrian Hamsters: Correlation with Offensive Aggression. Hormones and Behavior, 42(2), 182–191. 10.1006/hbeh.2002.1802

46. Jirikowski, G. F. (2019). Diversity of central oxytocinergic projections. Cell and Tissue Research, 375(1), 41–48. 10.1007/s00441-018-2960-5

47. Rokicki, J., Kaufmann, T., de Lange, A.-M. G., van der Meer, D., Bahrami, S., Sartorius, A. M., Haukvik, U. K., Steen, N. E., Schwarz, E., Stein, D. J., Nærland, T., Andreassen, O. A., Westlye, L. T., & Quintana, D. S. (2022). Oxytocin receptor expression patterns in the human brain across development. Neuropsychopharmacology, 47(8), 1550–1560. 10.1038/s41386-022-01305-5

48. Carstensen, L. L. (2006). The Influence of a Sense of Time on Human Development. Science, 312(5782), 1913–1915. 10.1126/science.1127488

49. Castle, E., Eisenberger, N. I., Seeman, T. E., Moons, W. G., Boggero, I. A., Grinblatt, M. S., & Taylor, S. E. (2012). Neural and behavioral bases of age differences in perceptions of trust. Proceedings of the National Academy of Sciences, 109(51), 20848–20852. 10.1073/pnas.1218518109

50. Ding, S.-L., Royall, J. J., Sunkin, S. M., Ng, L., Facer, B. A. C., Lesnar, P., Guillozet-Bongaarts, A., McMurray, B., Szafer, A., Dolbeare, T. A., Stevens, A., Tirrell, L., Benner, T., Caldejon, S., Dalley, R. A., Dee, N., Lau, C., Nyhus, J., Reding, M., … Lein, E. S. (2016). Comprehensive cellular-resolution atlas of the adult human brain. Journal of Comparative Neurology, 524(16), 3127–3481. 10.1002/cne.24080

