## Supplementary Material for "Oxytocin and vasopressin 1a receptor alterations in the superior temporal sulcus and hypothalamus in schizophrenia"

**Snowden et al. Supplementary Materials**

**Supplementary Table 1**

*Demographics for All Donors - Across Superior Temporal Sulcus and Hypothalamus Specimens*

| Donor ID | Group | Sex | Mean Age | Race |
|---|---|---|---|---|
| 3739 | Schizophrenia | Female | 62 | White |
| 5725 | Control | Female | 80 | Asian |
| 5744 | Control | Female | 77 | White |
| 5952 | Control | Female | 66 | Not reported |
| 6054 | Control | Male | 76 | White |
| 6227 | Control | Female | 47 | White |
| 5816 | Control | Female | 48 | White |
| 24578 | Schizophrenia | Male | 51 | White |
| 14502 | Schizophrenia | Male | 63 | White |
| 15957 | Schizophrenia | Male | 83 | White |
| 19459 | Control | Male | 63 | White |
| 26012 | Control | Male | 56 | White |
| 26914 | Schizophrenia | Male | 83 | White |
| 31998 | Schizophrenia | Male | 72 | Not reported |
| 34583 | Control | Male | 83 | White |
| 44271 | Schizophrenia | Female | 83 | White |
| 45611 | Schizophrenia | Male | 66 | White |
| 48185 | Schizophrenia | Male | 56 | White |
| 62199 | Schizophrenia | Male | 62 | Not reported |
| 63909 | Schizophrenia | Male | 62 | Not reported |
| 64175 | Schizophrenia | Female | 58 | White |
| 73787 | Control | Male | 66 | White |
| 76807 | Schizophrenia | Male | 76 | Asian |
| 79856 | Control | Female | 83 | White |
| 348862 | Control | Male | 82 | White |
| BEB18077A | Control | Male | 61 | White |

| Donor ID | Group | Sex | Mean Age | Race |
|---|---|---|---|---|
| HBCIA | Schizophrenia (paranoid) | Female | 58 | Black |
| HCT17HFQA | Control | Male | 59 | White |
| HCTYRA | Control | Female | 60 | White |
| S00281 | Schizophrenia | Female | 62 | White |
| S01119 | Schizophrenia | Female | 65 | White |
| S13800 | Schizophrenia | Female | 61 | White |
| S15221 | Schizophrenia | Female | 58 | White |
| S15348 | Schizophrenia | Female | 52 | White |
| S16818 | Schizophrenia | Female | 54 | Black |
| S13450 | Schizophrenia | Female | 64 | White |
| S17014 | Schizophrenia | Female | 56 | White |
| HCTZZJA | Control | Male | 70 | White |
| HCTZUA | Control | Male | 62 | White |
| 20475 | Schizophrenia | Male | 66 | Not reported |
| HCTZLA | Control | Female | 65 | White |

*Note.* Unless detailed otherwise, schizophrenia diagnoses were classified as schizophrenia (unspecified). All donors included in the SZ group had a confirmed diagnosis of SZ.

In general, diagnoses contained within the NBB inventory are classified based on the International Classification of Diseases (ICD-10) coding schema; however, for conditions that are inadequately represented by this categorization system, the NBB has generated codes and labels that best represent the condition(s) of the subject. For additional information on brain collection and neuropathology procedures, please see “NBB Best Practices” - https://neurobiobank.nih.gov/about-best-practices/

**Supplementary Table 2**

*Descriptive Statistics for Donors by Group and Sex – Hypothalamus OXTR Analysis*

| Group | Sex | Count | Mean Age | Standard Deviation |
|---|---|---|---|---|
| Schizophrenia | Female | 2 | 51.00 | 9.9 |
| | Male | 5 | 71.80 | 12.6 |
| Control | Female | 3 | 57.67 | 9.71 |
| | Male | 4 | 63.00 | 4.83 |

*Note.* Total of 14 donors included in the receptor density analysis after exclusion of 14 donors due to anatomical variation and a lack of signal available to quantify.

**Supplementary Table 3**

*Descriptive Statistics for Donors by Group and Sex – Hypothalamus AVPR1a Analysis*

| Group | Sex | Count | Mean Age | Standard Deviation |
|---|---|---|---|---|
| Schizophrenia | Female | 1 | 44.00 | NA |
| | Male | 6 | 71.83 | 11.27 |
| Control | Female | 1 | 47.00 | NA |
| | Male | 4 | 64.75 | 4.11 |

*Note.* Total of 12 donors included in the receptor density analysis after exclusion of 16 donors due to anatomical variation and a lack of signal available to quantify.

**Supplementary Table 4**

*Descriptive Statistics for Donors by Group and Sex – Superior Temporal Sulcus OXTR Analysis*

| Group | Sex | Count | Mean Age | Standard Deviation |
|---|---|---|---|---|
| Schizophrenia | Female | 11 | 60.45 | 8.77 |
| | Male | 11 | 68.18 | 9.26 |
| Control | Female | 6 | 69.67 | 13.32 |
| | Male | 9 | 66.89 | 9.10 |

*Note.* Total of 37 donors included in the receptor density analysis after exclusion of 4 donors due to film artifacts or tissue damage.

**Supplementary Table 5**

*Descriptive Statistics for Donors by Group and Sex – Superior Temporal Sulcus AVPR1a Analysis*

| Group | Sex | Count | Mean Age | Standard Deviation |
|---|---|---|---|---|
| Schizophrenia | Female | 10 | 61.50 | 8.38 |
| | Male | 10 | 68.90 | 9.26 |
| Control | Female | 6 | 68.83 | 13.81 |
| | Male | 10 | 67.80 | 9.59 |

*Note.* Total of 36 donors included in the receptor density analysis after exclusion of 5 donors due to film artifacts or tissue damage.

**Supplementary Table 6**

*Donor 44271 Brainbank Information*

| Brain Weight (g) | pH | Age at Date of Collection | Postmortem Interval (hours) |
|---|---|---|---|
| 1186 | 6.07 | 83 | 26.75 |

**Supplementary Figure 1**

*Autoradiogram Showing AVPR1a Binding in the Superior Temporal Sulcus for Donor 44271*

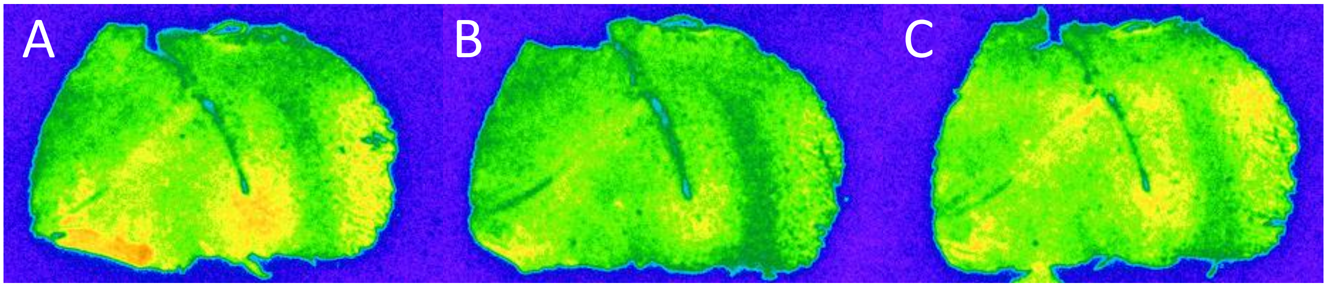


*Note*. A) AVPR1a radioligand alone. B) AVPR1a radioligand with an AVPR1a antagonist C) AVPR1a radioligand with an OXTR antagonist

**Supplementary Figure 2.** Correlation between Donor Chlorpromazine Equivalent Dosage and Mean AVPR1a Densities in the Superior Temporal Sulcus

*Note*: Bands represent +/- 1 standard error of the mean. Medication records from 8 donors were available from specimens received from the Mt. Sinai NeuroBioBank.

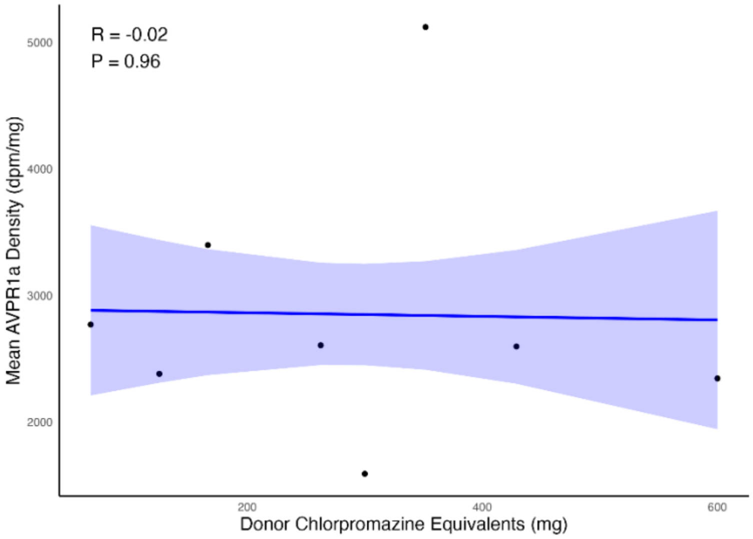


**Supplementary Figure 3.** Correlation between Donor Chlorpromazine Equivalent Dosage and Mean OXTR Densities in the Superior Temporal Sulcus

*Note*: Bands represent +/- 1 standard error of the mean. Medication records from 8 donors were available from specimens received from the Mt. Sinai NeuroBioBank.

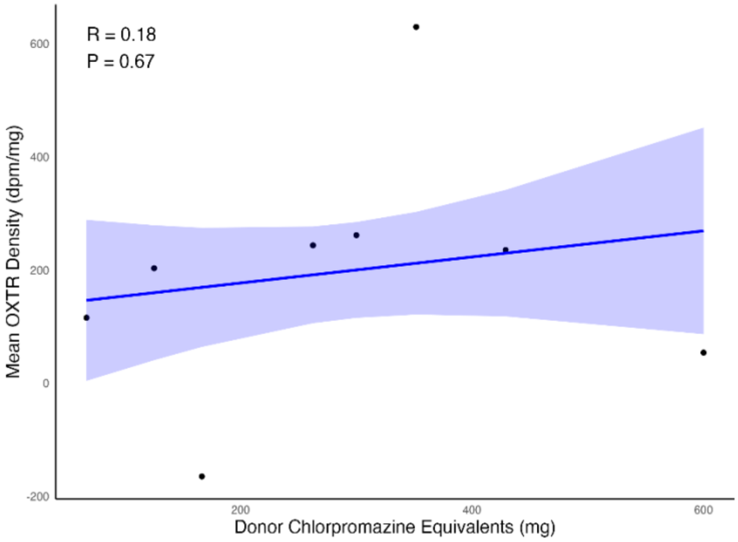

**Supplementary Figure 4**

*Autoradiograms Showing AVPR1a Radioligand Binding in the Hypothalamus*

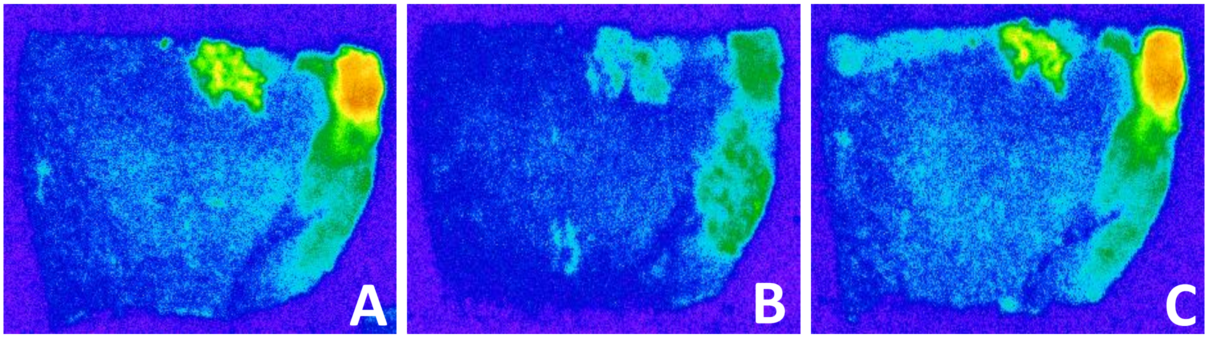


*Note.* A) AVPR1a radioligand alone. B) AVPR1a radioligand with an AVPR1a antagonist C) AVPR1a radioligand with an OXTR antagonist. Panel A shows the combined AVPR1a and OXTR receptor signals. Panel B shows the OXTR signal, as a result of the AVPR1a signal being eliminated by the AVPR1a antagonist. Panel C shows the isolated AVPR1a signal.

**Supplementary Figure 5**

*Autoradiograms Showing OXTR Radioligand Binding in the Hypothalamus*

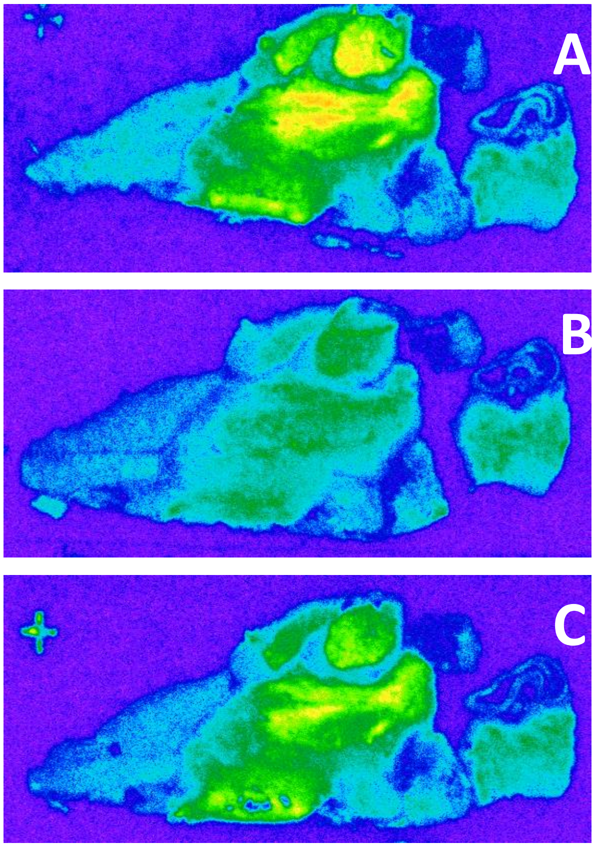


*Note***.** A) OXTR radioligand alone. B) OXTR radioligand with an OXTR antagonist C) OXTR radioligand with an AVPR1a antagonist. Panel A shows the combined OXTR and AVPR1a receptor signal. Panel B shows the AVPR1a signal, as a result of the OXTR signal being eliminated by the AVPR1a antagonist. Panel C shows the isolated OXTR signal.
